# Assessment of Augmented Reality Glasses for Spatial Tracking and Intraoperative Annotation in Veterinary Surgery

**DOI:** 10.64898/2025.12.18.695281

**Authors:** Yash Tipirneni, Andrew Blandino, Boaz Arzi, Chrisoula Agape-Toupadakis Skourita, Stephanie Goldschmidt

**Affiliations:** University of California – Davis, Department of Biomedical Engineering, Davis, CA, USA; University of California – Davis, Department of Statistics, Davis, CA, USA; University of California – Davis, School of Veterinary Medicine, Department of Veterinary Surgical & Radiologic Sciences, Davis, CA, USA

**Keywords:** augmented reality, hand-tracking software, intraoperative navigation, 3D hologram, medical technology

## Abstract

**Objectives:** Augmented reality (AR) glasses may improve surgical precision by projecting holographic overlays directly onto the surgical field. This study aimed to evaluate the feasibility of AR technology for enhancing spatial tracking.

**Methods:** We developed an AR application in Unity compatible with XReal glasses that allowed users to annotate and interact with a realistic 3D hologram of a dog head. Resident and specialist veterinarians were recruited to completed coordinate (distance) and outline (area) annotations under two conditions: (1) transfer: memorizing targets from a computer screen, and (2) direct: seeing the targets directly on the head. Distance errors, area metrics, and completion times were recorded from each participant.

**Results:** The mean distance error (N = 22) was significantly lower for direct versus transfer coordinates (2.73 ± 0.79 mm vs. 3.42 ± 1.81 mm). Area coverage (N = 20) was higher (83.7% ± 13.4% vs. 63.3% ± 16.2) and non-overlap was similarly reduced with AR-guidance. Completion times differed significantly between the transfer and direct groups for coordinate tasks (11.2 ± 10.4 sec versus 8.19 ± sec) but not for area tracing (25.7 ± 18.3 sec versus 26.8 ± 26.5 sec).

**Conclusion:** AR-guided visualization improved spatial accuracy for both distance and area metrics without reducing speed. The effects observed for specialty, eyewear, or arm length were negligible. However, the level of experience with a cutoff of 2 years did have a significant effect on distance error.

**Clinical Relevance:** These findings support the utility of AR for optimizing surgical precision in veterinary medicine.

## Introduction

In surgical oncology procedures, the accurate delineation of tumor margins is critical to ensure the complete removal of cancer cells.^1,2^ Yet, there is currently no accepted intraoperative tool for accurate delineation of cancer versus normal tissue. Thus, a surgical safety margin (removal of grossly normal surrounding tissue) is often applied to remove all non-visible cancer cells. When a wide surgical margin is used, the chance of local recurrence is minimized, but the patient’s morbidity increases.^3^ Consequently, if the surgical margins are underestimated, the likelihood of tumor recurrence or persistence and diminished survival times will increase.^4–6^ Therefore, accurately determining cancer from normal tissue remains a critical challenge within clinical practice. To address these challenges, multiple new intraoperative tools are being clinically investigated.

Computer-aided intraoperative systems are one such tool that may empower clinicians to make informed decisions during procedures. They acquire pathological and spatial data from biological tissues^7–10^ and display the sensor output on a monitor, often accompanied by a camera feed. However, when image overlays are displayed on top of a 2-dimensional (2D) recording, the video file lacks binocular depth perception and 3-dimensional (3D) spatial orientation.^11,12^ Surgeons must then manually translate these 2D coordinates into 3D space, often approximating the precise location on body structures. This menial process not only diverts attention away from the surgical field but can also result in sub-centimeter inaccuracies.^13,14^

To overcome such limitations, human head and neck surgeons are in a process of adapting 3D surgical planning approaches to enhance point-to-point tracking.^15,16^ By aligning pre-operative annotation holograms with anesthetized patients, otolaryngologists have been able to reduce spatial uncertainty, thereby improving outcomes.^15^ One method for displaying sensor data or anatomical structures directly in the field of view^17^ is through augmented reality (AR) glasses with a birdbath optical design.^18^ In this setup, a micro-OLED display projects an image onto a beam splitter, which then reflects the light toward a concave mirror. The mirror then focuses the rays directly into the user’s eye, allowing holographic images to appear seamlessly within real-world environments. While this technology is becoming increasingly common with physicians,^19^ the utilization of AR technologies by veterinary surgeons is in its early stages.^20^

The aim of this study was to evaluate the practicality of AR glasses in enhancing the tracking of annotations using an oral tumor in the dog as a model . To assess feasibility, participants completed distance and area accuracy tasks on a cadaveric dog hologram. This work represents an initial step toward integrating augmented reality into veterinary surgical oncology, with the potential to improve operative precision and patient outcomes.

## Methods

The Air 2 Ultra^21^ AR glasses (XReal [formerly NReal]; Beijing, China) were utilized in this study. The glasses have a birdbath design that is capable of projecting holograms directly into the wearer’s field of view. The glasses were connected to the Beam Pro console (XReal [formerly NReal]; Beijing, China) via a high-speed data transfer USB-C cable. The Beam Pro is an Android-based device specifically designed by XReal for their glasses that functions as both a controller and a spatial-computing hub. Together, they enable hand tracking and fingertip detection, which are integral to the annotation component of this study (Figure 1).

**Figure 1:**
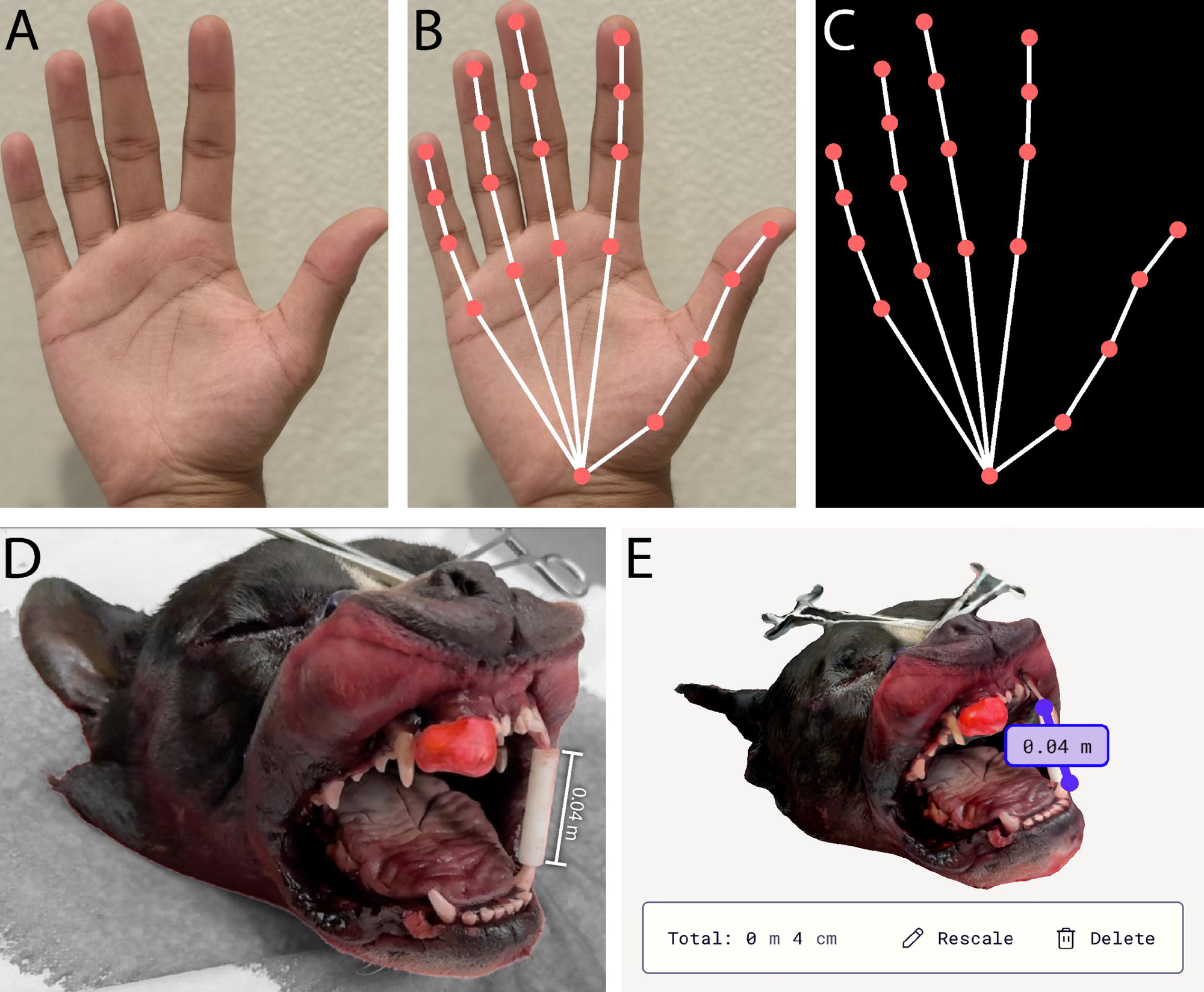
(A) Photograph of a person’s palm representing what a user will see without the AR glasses (B) A person’s palm is overlaid with vector-line representations generated from Python code. This is what the user will see while wearing the glasses (C) The vector-line representation is displayed on a black background without the person’s palm. This is what a screenshot or video will register from the augmented reality (AR) glasses. (D) Photograph of a cadaver dog head with a clay mold fitted around its right maxillary incisors to model an oral tumor. (E) The 3D reconstruction scan was created by the Polycam app after taking 124 images in Light Detection and Ranging (LIDAR) mode. The ruler tool shows that the mouth gag is 4 centimeters (cm) long, confirming the 3D object is a 1:1 scale replica.

The AR application was developed using Unity,^22^ with custom scripting in C#. Unity provided the core framework for creating interactive 3D GameObjects, while C# scripts-controlled features such as panel navigation, model rotation, and annotations. The interface was designed with a simple panel that allows users to select functions, capture images, and access supporting documentation by simply pressing buttons. The application was integrated with the XReal SDK to render holographic overlays directly through AR glasses. Hand tracking was enabled using the Android XR plugin, allowing users to interact naturally with virtual models without controllers, while vector-line representations were displayed directly on their fingers. Colliders detected fingertip interactions, enabling precise annotation of the hologram. This setup provided an intuitive interface while maintaining the accuracy required for anatomical assessment.

To support surgical evaluation, the app included interactive, user-guided programs specifically designed to assess spatial accuracy. The guided program featured custom algorithms that were implemented to calculate both distances and area transference. Inter-point distances were computed using the standard Euclidean metric in Unity’s native system, with results in millimeters. Since the object was a 1:1 replica of the cadaver, using a scale factor was not necessary.

The program estimates the area of the region of interest (ROI) by projecting the fingertip trace-path onto a 2D plane as a flattened polygon. It uses the shoelace formula^23^ to compute the enclosed region in squared millimeters. Both the user-drawn outline and the target region are parsed in this fashion for consistency. The app then compares the overlap between the plotted and target regions using pixel-based interpolations. This approach improves the accuracy of coverage estimation even on anatomically complex, curvilinear surfaces.

We acquired a canine cadaver head from animal that was euthanized for reasons unrelated to this study and custom fitted it with clay (CiaraQ; Shenzhen, China) mold covered with acrylic paint (Crafts4All; Boksburg, South Africa) and gloss varnish (Liquitex; Piscataway, NJ) to stimulate a rostral maxillary gingival tumor. Multiple lip retractors (IM3; Vancouver, WA) were used to retract the lower and upper lips, and a mouth gag (IM3; Vancouver, WA) was placed between the maxillary and mandibular canine teeth to stimulate a clinical imaging situation. Further, two studio lights (Torjim; Shenzhen, China) were positioned opposite from each other to minimize shadows during filming and simulate the brightness of operating room lights.

An iPhone 16 Pro (Apple Inc.; Cupertino, CA), which has a Light Detection and Ranging (LIDAR) scanner, was mounted to a gyroscopic camera stabilizer (Aochuan; Shenzhen, China). LIDAR technology is capable of estimating distances with notable accuracy and is commonly used for precise 3D reconstruction.^24^ The Polycam app^25^ was utilized to create a 1:1 3D scans of the dog heads with the built-in LIDAR mode.^26^ The scans were later cropped to the exact dimensions of the heads and exported as .OBJ files (Figure 1).

### Participant Testing

Participants (residents and faculty) were recruited from the dentistry and oral surgery, neurology and neurosurgery, and orthopedic surgery services at the UC Davis School of Veterinary Medicine. Prior to participant recruitment, approval was obtained from the University of California, Davis Institutional Review Board (IRB #2333805) under an exempt status.

Participants were allowed 2 minutes to practice drawing on the dog head hologram to become acquainted with the software before we collected data. For the distance testing, participants were shown 5 different coordinates on a computer screen with an image of a cadaver head and had 2.5 seconds to memorize the locations. They were then instructed to transfer those points to the hologram. Afterwards, the coordinates were displayed directly on the hologram, and they had to draw a point at the center. Time taken to complete the tasks and the distance offset from the target coordinate in millimeters were recorded for both parts.

Then, the participants were shown a region of interest (ROI) outline on an image of the head on a computer screen and had 5 seconds to memorize it. They were similarly told to transfer the ROI to the hologram. After the ROI was displayed on the hologram, they had to trace the outline. Two different area metrics were recorded: the area of non-overlap and the ROI-coverage. The area of non-overlap was defined as the area that was drawn that fell outside the ROI. The ROI coverage was the area of the ROI that was covered by the user’s drawing. The time to draw each ROI was recorded as well (Figure 2).

**Figure 2:**
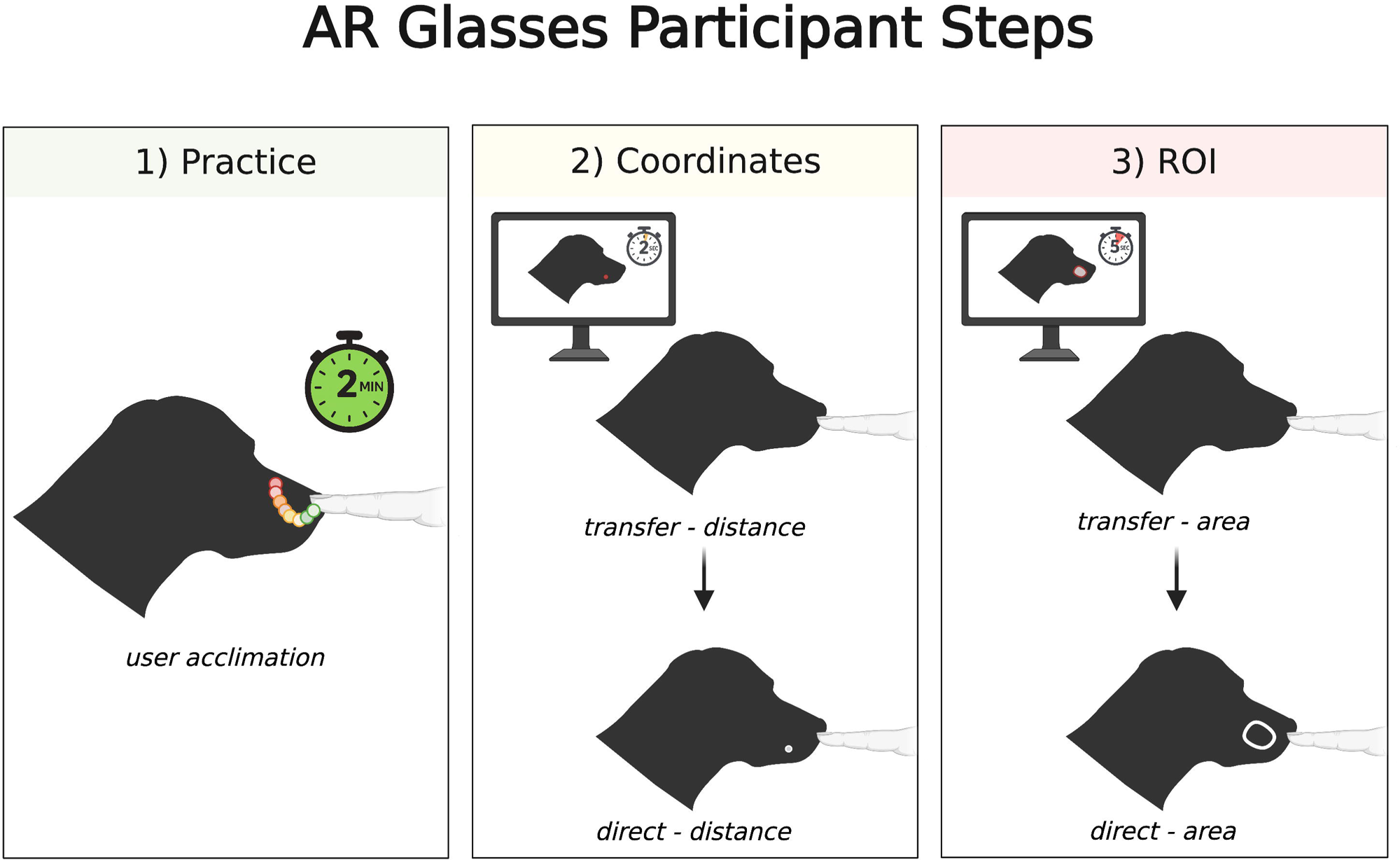
Visual representation of how spatial accuracy was evaluated with the AR goggles. (1) Each participant has 2 minutes to practice drawing to become acclimated to the program. (2) The participants will then transfer 5 coordinates from a computer screen to the hologram from memory, and then the coordinates will appear directly on the hologram and be directly marked by the user. (3) The participants will transfer 3 regions of interest (ROIs) from a computer screen to the hologram from memory, and then the ROIs will appear directly on the hologram for the user to trace directly. The area of coverage and overlap for each trial was recorded in squared millimeters.

The purpose of the 2D to 3D “transfer” tasks was to obtain a raw metric related to human error from memory and cognition. The “direct” tasks, where the user drew a point at the center of a coordinate or traced an ROI, were performed as a baseline measurement to assess various factors, such as dexterity, depth perception, and the instrumental bias of the hand detection software. The transfer data was normalized by the direct data to estimate the error of the 2D to 3D transference created while also accounting for these confounding variables.

### Statistical Analysis

Continuous outcomes were reported as mean ± standard deviation or median (IQR) depending on the data distribution. Data distributions were evaluated for normality to choose between parametric and non-parametric testing. Parametric tests were used when normality and homogeneity of variance were met. Other statistical tests were selected according to data characteristics and the study design. Paired t-tests were used to compare repeated participant distance and area data for transfer vs direct conditions. Variance among the 3 specialties was evaluated using one-way ANOVA (parametric) and Kruskal-Wallis (non-parametric) tests.

Independent group comparisons of demographic data were conducted with Welch’s t-tests (parametric and unequal variance) and Mann-Whitney U-tests (non-parametric). A standard linear regression was performed to compare participant arm lengths with their respective distance error. The tests were 2-tailed to allow detection of effects in either direction. All statistical analyses were carried out with Microsoft Excel and Python (Google Colab), using the *pandas*, *scipy.stats*, *statsmodels*, and *numpy* libraries. Statistical significance was defined as p < 0.05.

## Results

We enrolled 22 licensed veterinarians from the UC Davis School of Veterinary Medicine in the study. Of these veterinarians, 7 were from the dentistry and oral Surgery Service, 7 were from the orthopedic surgery Service, and 8 were from the neurology and neurosurgery Service. The study group consisted of 3 first-year residents, 7 second-year residents, 3 third-year residents, 1 fellow, and 7 board-certified faculty members. Only one veterinarian was left-handed, while the rest were right-handed.

### Spatial Accuracy

The software accurately calculated the distance error of points placed on the dog head hologram. The spatial accuracy tasks (Figure 3) had different performance metrics between the direct and transfer conditions (Figure 3A). The average distance error for the transfer coordinates was 3.42 ± 1.81 mm, which was significantly higher (p = 2.36E-4; t = 3.60; df = 21) than the distance error for the direct coordinates, which was 2.73 ± 0.79 mm (Figure 3B). The mean normalized difference (transfer - direct) across users was 0.68 ± 1.98 mm, with a small to moderate effect size (Cohen’s d = 0.34). Similarly, the average task completion time (Figure 3C) was significantly (p = 1.82E-3; t = 2.97; df = 21; d = 0.28) greater in the transfer group (11.2 ± 10.42 seconds) than in the direct group (8.19 ± 5.36 seconds). These results demonstrate that visualizing coordinates in one’s field of view improved both precision and efficiency in terms of distance errors.

**Figure 3:**
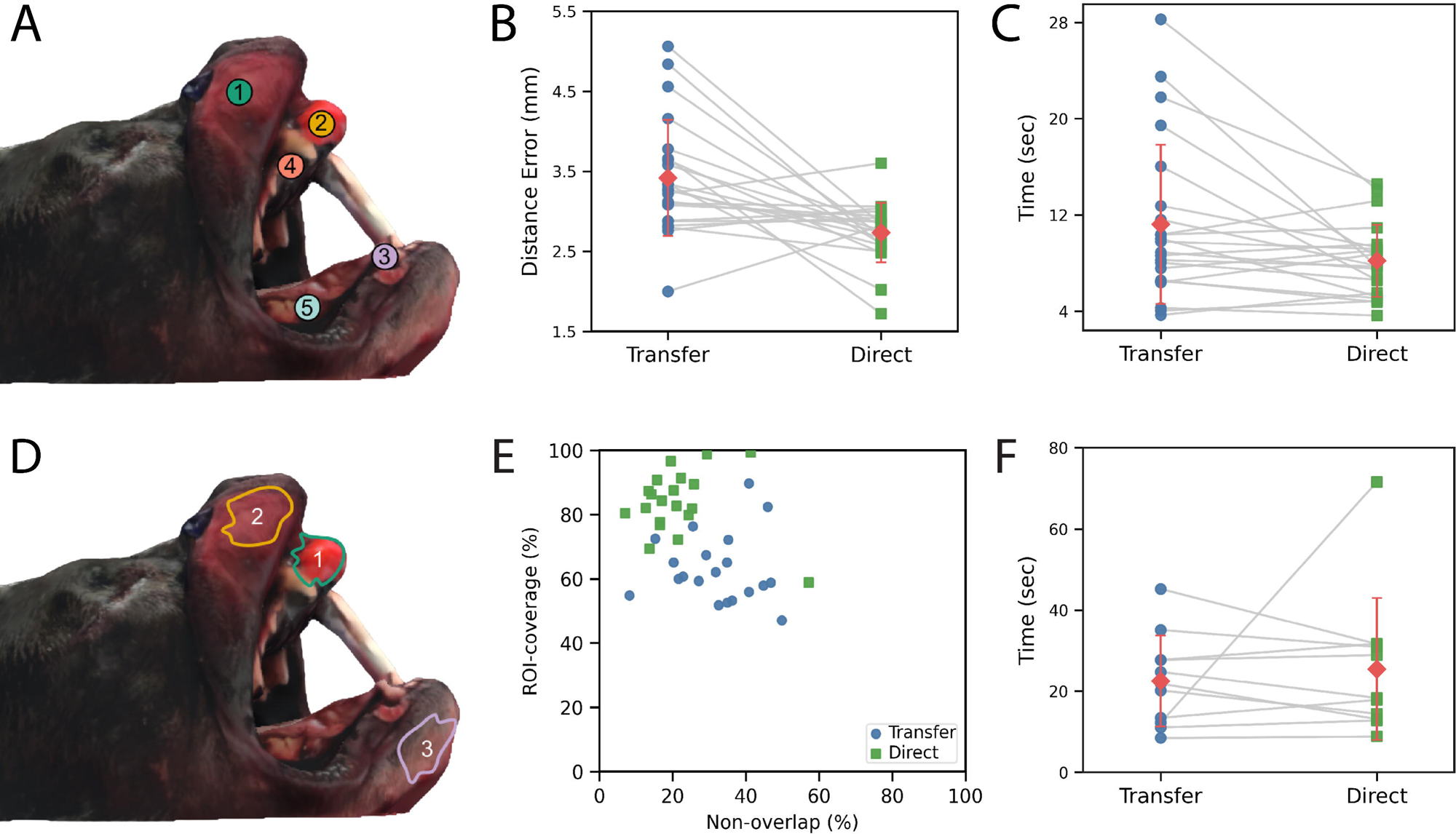
(A) A map of the location of the five coordinates on the dog head hologram. (B) A comparison of the transfer (estimating coordinate location on the hologram from viewing the markers on a computer screen) and direct (tapping a coordinate directly on the hologram) results for each participant’s average distance error in millimeters. (C) A comparison of the average time each participant took for the transfer and direct groups. (D) The location of the 3 regions of interest (ROIs) on the virtual dog head. The data from each participant’s 3 runs are averaged for statistical simplicity. (E) Comparison of the non-overlap percentage (percentage of the user-drawn ROI that falls outside the target ROI) with the ROI-coverage percentage (percentage of the target ROI that is covered by the user-drawn ROI). The direct (target is shown directly on the hologram) and transfer groups (ROI is shown on a computer screen and then needs to be drawn on the hologram) are also shown separately. (F) The time taken to draw an ROI is shown for each participant for the transfer and direct groups.

The software was able to calculate the region of interest (ROI) areas accurately for 3 pre-programmed boundaries (Figure 3D); the accuracy area estimations were qualitatively verified by testing the algorithm on various planar and freeform surfaces, and the outputs were consistent with visually expected values. On average, the user-drawn contours covered the majority (> 50%) of the ROIs, and the area of overlap generally exceeded the area of non-overlap (Figure 3E). Overall, the difference between transfer and direct contours was statistically significant (Figure 6B) for both the ROI area coverage (p = 2.23E-12) and area of non-overlap (p = 2.15E-04) but insignificant for the time taken (p = 0.397; Figure 3F). The average area of the transfer group was 63.3% ± 16.2% and 32.2% ± 19.9% whereas the direct group was 83.7% ± 13.4% and 21.7% ± 14.1% for the ROI area coverage and the area of non-overlap, respectively. The data shows that an augmented reality (AR) overlay may help better pinpoint target tissue while minimizing the misidentification of adjacent, non-target tissues.

### Demographics

Figure 4 depicts how user demographics (N = 22) impact spatial accuracy. The median (IQR) averaged, normalized errors for orthopedic surgery (N = 7), neurosurgery (N = 8), and oral surgery (N = 7) were 0.580 (0.030 - 0.650) mm, 0.280 (0.105 - 1.200) mm, and 0.944 (0.350 - 2.080) mm, respectively (Figure 4A). The distance error did not differ significantly between specialties (one-way ANOVA; p = 0.337). A non-parametric test confirmed this (Kruskal-Wallis; p = 0.459), indicating no group-level effect. The clinicians’ years of experience (Figure 4B) were significant between those with ‘≤ 2’ (median = 0.990 years; IQR = [0.180, 1.785]; N = 14) and ‘> 2’ (median = 0.070 years; IQR = [-0.305, 0.610]) years based on a Welch’s t-test (t = 2.845; p = 0.0104) and a Mann-Whitney U-test (U = 90.0; p = 0.0197).

**Figure 4:**
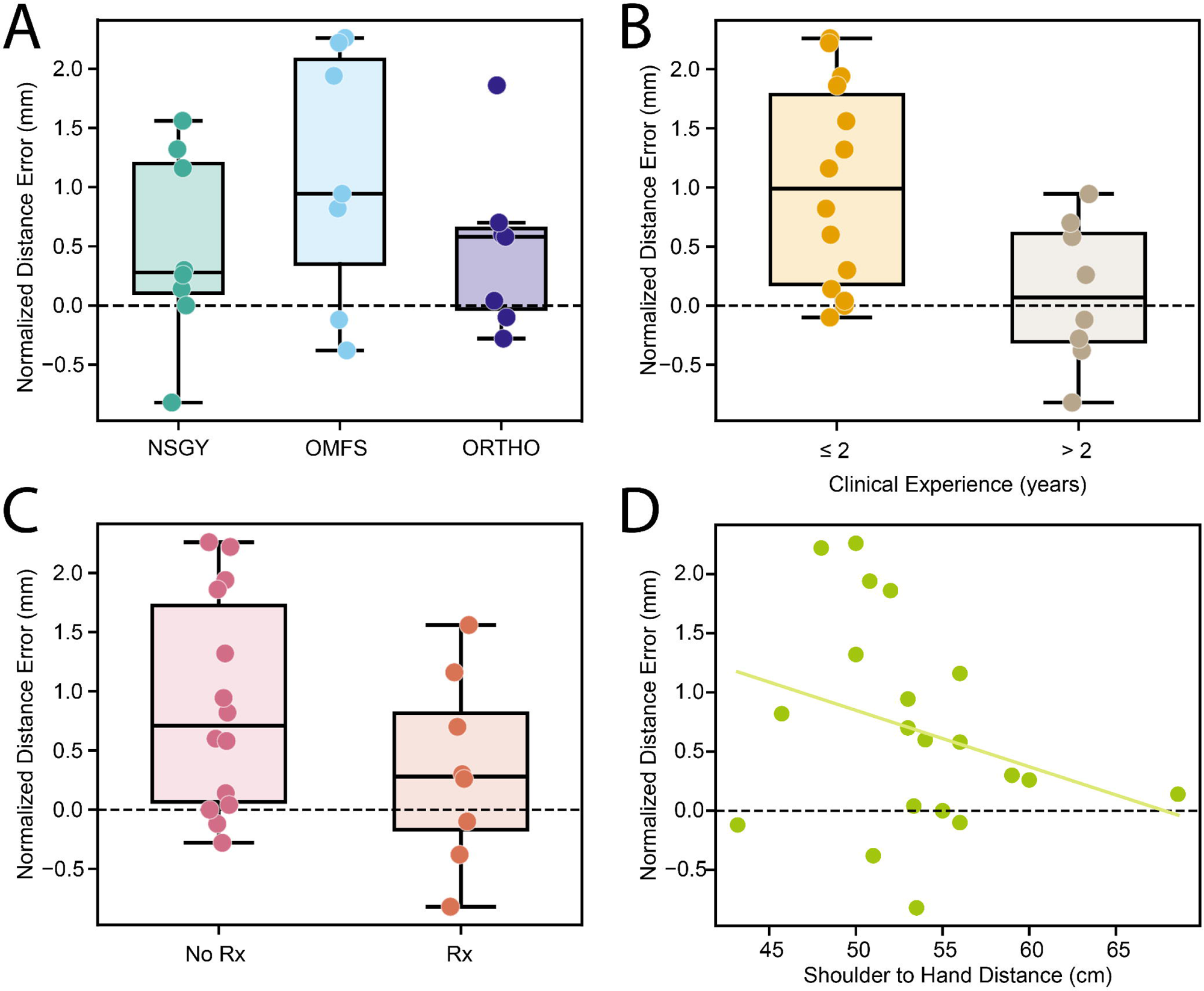
Comparison of various demographics to the average normalized distance error in millimeters. (A) Plot comparing the median error for the 3 specialties (NSGY = neurosurgery; OMFS = oral & maxillofacial surgery; ORTHO = orthopedic surgery). (B) Plot comparing the median error for participants with less than or greater than 2 years of clinical experience. (C) Plot comparing the median error of participants with and without optical prescriptions. (D) Plot comparing the participants’ arm length to their respective errors. The trendline has a negative slope.

The median (IQR) averaged, normalized error was 0.710 (0.065 - 1.725; N = 14) mm for participants without prescription eyewear and 0.280 (−0.170 - 0.815; N = 8) for participants with prescription eyewear (Figure 4C). The difference between the groups was insignificant based on a Mann–Whitney U-test (U = 74.0; p = 0.238). There was also a poor correlation between one’s arm length (N = 20) and their performance (Figure 4D); a linear regression model had the formula, -0.0477x + 3.2316 (R^2^ = 0.0864; p = 0.208).

### User impression

Two participants did not complete the ROI tests and the survey. The median (IQR) years of experience practicing medicine was 2.0 (1.0-8) years, with a range of 0-26 years. None of these participants were wearing contact lenses at any point and did not use optical inserts in the AR glasses. The median (IQR) optical prescription for participants with glasses (N = 7) was - 2.00 (−4.94 to -0.813) diopters (D) when their spherical equivalents were averaged between eyes, with a range of -7.25 to 4.50 D. The participants’ median (IQR) distance (N = 20) between their shoulder and wrist with their hand fully extended was 52.5 (42.75-55.25) centimeters (cm), with a range of 17 to 60 cm. The average participant satisfaction score (Figure 5) across the 10 survey questions was 3.55 ± 0.934 on a scale of 1-5. The lowest (natural/responsive) average for a question was 2.75 ± 0.911 out of 5, while the highest (intuitive) was 4.10 ± 0.641.

**Figure 5:**
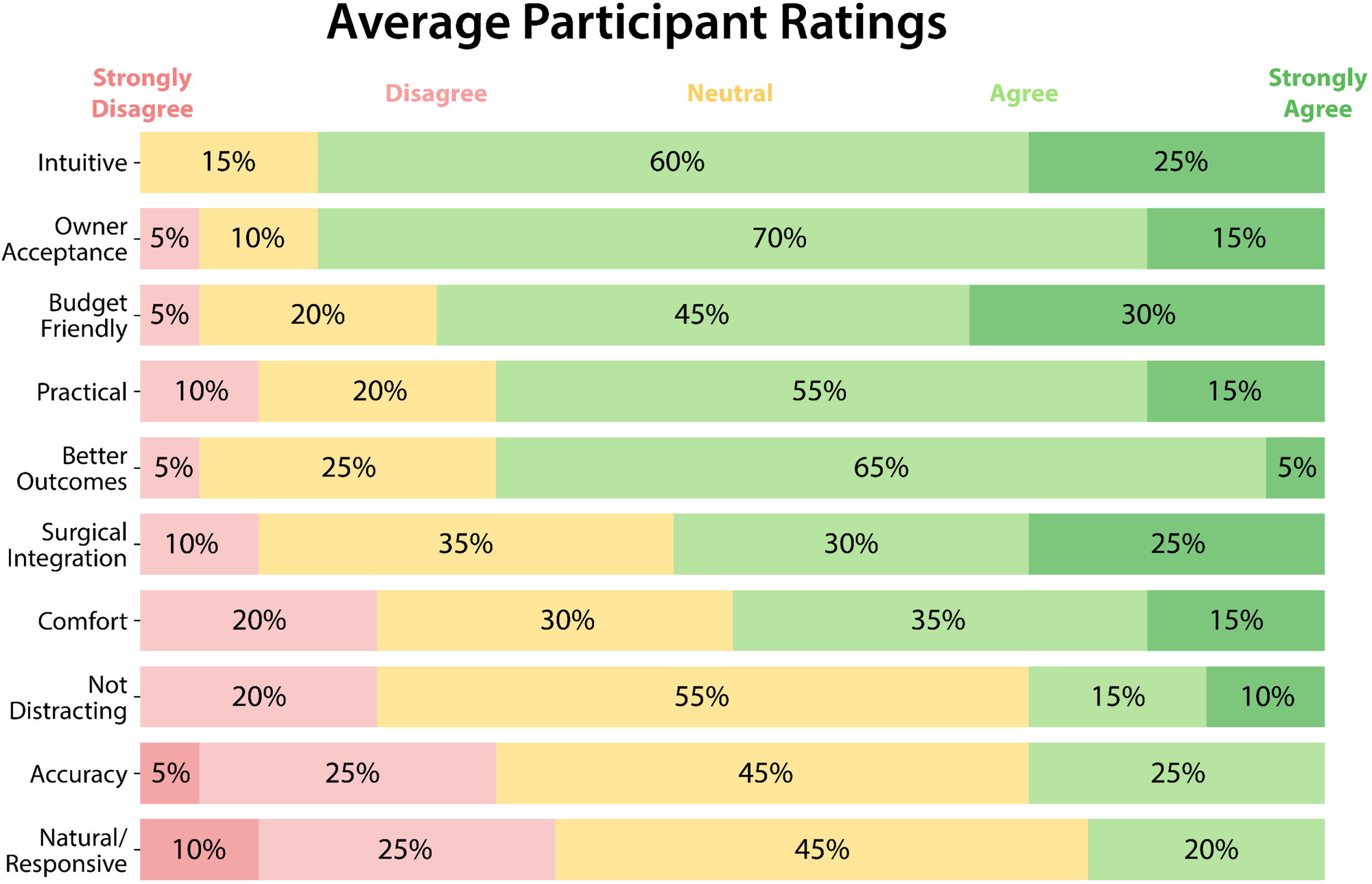
Qualitative ratings provided by the study participants, in which they rated 10 different aspects of the AR technology on a scale of 1 to 5, with 1 being strongly disagree and 5 being strongly agree.

## Discussion

The data from this study indicated that augmented reality (AR) tools may enhance spatial tracking accuracy for oral tumor within the context of the oral and maxillofacial surgical (OMFS) anatomy. Participants achieved lower normalized errors with AR for both point-to-point measurements and the region of interest (ROI) mapping. Moreover, the coordinate tasks with AR-guidance were significantly faster than the standard screen-to-field transfer method, suggesting that AR may improve surgical workflow. Consequently, the time taken for the AR ROI tasks was negligible between the AR-guidance and the transfer datasets. This implies that AR may improve user accuracy across various tasks while not compromising on speed or efficiency.

The improved accuracy with AR is likely related to its ability to mitigate the 3D spatial uncertainties commonly associated with monitor-based image-guided surgery setups.^27,28^ In standard practice, surgeons view a monitor screen and must mentally translate spatial and pathological data onto a 3D patient, which is a process prone to sub-centimeter errors. By superimposing holographic annotations directly onto the surgical field, AR technologies allow surgeons to visualize ROIs in situ without looking away.^29^ This direct integration eliminates hypothetical guesswork in the translation process, thereby reducing spatial error. High precision is imperative for clinical applications, such as surgical oncology, where even slight margin errors can risk removing healthy tissue or leaving cancer cells behind.^1,2^ Thus, AR may become essential in computer-aided surgery such as optimizing tumor margin delineation intraoperatively (Figure 6).

**Figure 6:**
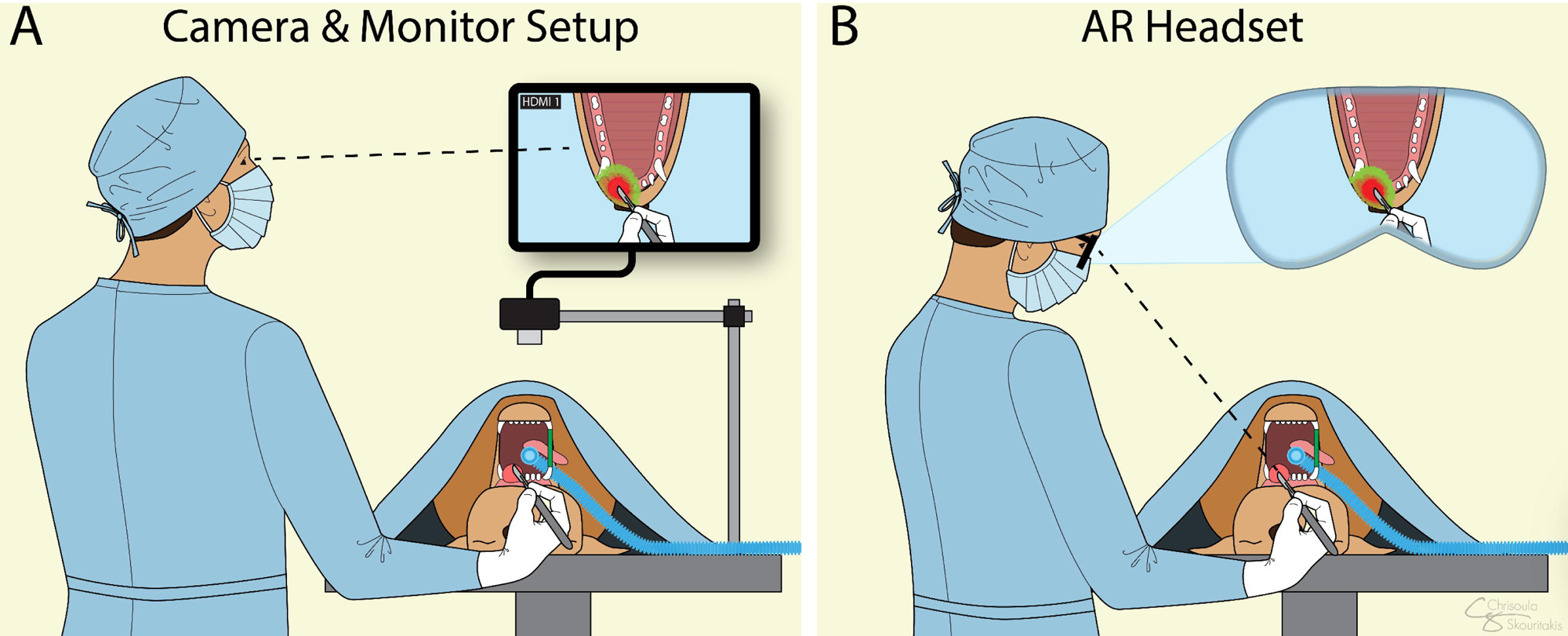
Concept figure of how this augmented reality (AR) technology may improve clinical practice. (A) The surgeon looks at a computer screen while using the tumor mask overlay to inform their surgical margins. (B) The surgeon looks directly at the patient while the mask overlays are projected into their field of view as AR holograms.

Surprisingly, direct AR mapping did not significantly reduce the time taken for outline annotations. While outlining an irregular tumor area is inherently time-consuming, participants may have dedicated more time to precisely trace the ROI when it was visible directly on the hologram as opposed to when they saw it on a computer screen for only 5 seconds. It may be possible that any time saved by not switching focus between the computer screen and dog head was offset by the effort required to maneuver AR controls or to maintain hand stability mid-air. The participants’ learning curves or system latency could have also affected task metrics.

Nevertheless, while AR did not make area tracing faster, it made it more precise. In surgical oncology, precision is considered more important than speed since patient outcomes are dependent on tumor clearance. Thus, AR devices may deliver better outcomes during surgeries and may prove to be a strong asset in the future.

Beyond distance errors, several participant demographic trends became apparent. For example, neurosurgeons slightly outperformed OMFS participants in spatial tasks. This may reflect individual variations rather than true specialty-level performance. Both specialties operate in confined, anatomically complex spaces.^30^ However, the small sample size of 7-8 participants per group limits us from drawing definitive conclusions. Further research with larger cohorts would be beneficial to explore whether a difference in surgical training influences one’s performance in AR-guided workflow. Another trend was that participants with more experience tended to achieve better accuracy, however the weak correlation and small sample size caution against over-interpretation. The older clinicians likely had more surgical experience, which may have helped them perform better on the spatial accuracy tests. This experiential effect, while not robust, aligns with the expectations that senior clinicians may confer an advantage in surgical simulation tasks.^31^

It was also noted that participants who used optical prescriptions unexpectedly outperformed those without optical prescriptions on the coordinate tasks. One possible explanation is that these individuals were all nearsighted and could naturally focus on close-up holographic details^32^ better than those with perfect distance vision. Given the small sample size, the results should be interpreted cautiously, but this raises an important point on how visual acuity impacts AR experience. There was an inverse, albeit poor, correlation between arm length and distance error. This may be because hand-tracking is more challenging with the external camera on the AR glasses when the user’s arms are too short. Moreover, each specialty group was tested in different rooms, so minor differences in ambient lighting may have played a role. Overall, these subgroup observations may be a promising area of study for future medical AR research.

The skin tone of the participants may have played a small factor during testing since the glasses’ camera was pointed at the back of their ungloved hands. However, during app development and testing, we noticed that hand-tracking performed well on both light and dark skin-tones. The skin colors of participants were not recorded during the study, but we did not notice any correlation with performance. The races/ethnicities in this study were majority white but also included Asian, Hispanic/Latino, and Black people. We did not ask participants to disclose their genders, but from our interactions, we infer that approximately ⅔ of the clinicians identified as women. No one self-reported any disabilities that would impact their abilities to perform surgery. Additionally, other factors such as the facial fit of the AR glasses or the user’s ergonomics could have marginally impacted hand-tracking depending on how far one’s hands deviated from the external camera viewport during the software runtime.

Several limitations of this study should be considered. Foremost, the sample size (20-22) was small, and participants were all recruited from a single department within one institution, which may limit generalizability. The trends noted could be amplified or perhaps minimized in a larger and more diverse cohort. Additionally, the speed of user tasks was recorded manually, which may have introduced minor errors. Also, the experimental tasks were performed on a holographic simulation rather than on a physical cadaver. While this approach enabled the accurate quantification of spatial metrics, real surgical conditions (i.e., bodily fluids, intensive lighting, multiple people) would present additional challenges that were not accounted for. The actual feasibility of AR-integration into veterinary surgical oncology is yet to be determined, but this pilot study was an important first step.

The AR system also presented additional testing constraints. When the app was running for extended periods (> 90 minutes), we observed some lagging and decreased responsiveness in hand-tracking, likely due to the processing limits of the Snapdragon 6 Gen 1 chip (Qualcomm; San Diego, CA) in XReal’s Beam Pro device. Each set of participant specialties was tested on 3 different dates to allow for system resets, which minimized performance degradation. Still, this highlighted how current commercial AR glasses struggle with prolonged, continuous usage.

More robust AR headsets, like the HoloLens 2 (Microsoft; Redmond, WA), have more powerful processors and may ensure smoother performance during longer sessions. In addition, the learning curve is important consideration as although all participants were given 2 minutes to acclimate to the application, their performance may have improved with more sessions. Despite these limitations, this study demonstrated the feasibility and benefit of AR-integration for surgical planning. The insights gained here may inform larger, follow-up studies and eventually, clinical trials. Future work should focus on refining the software to minimize lag, improving hand-tracking, and accounting for the tissue manipulation in real patient surgeries.

Overall, this investigation demonstrates that AR-integration may prove useful in surgical precision for veterinary surgery. The ability to project tumor outlines directly onto the surgical field appears to confer greater accuracy without sacrificing efficiency over the conventional monitor-based approach. Taken together, the findings support AR-guidance as a promising adjunct for image-guided surgeries that require high spatial accuracy, offering an improvement over currently used paradigms. As AR technologies become more prevalent in medical settings and user-friendly, the barriers to adoption may inevitably diminish. Widespread acceptance of AR surgical guidance therefore may be on the horizon. A new systematic review^20^ on augmented and virtual reality (AR/VR) devices within veterinary medicine specifically highlighted this significant gap in surgical care that remains unaddressed. Our study, therefore, lays the groundwork for future exploration of AR and its potential within veterinary surgery.

## Acknowledgments

We would like to thank the clinicians who took the time to participate, the UC Davis SVM for sponsoring this study, and the owners who donated their dog’s bodies.

## Disclosures

### AI-assisted technology

Copilot in Visual Studio Code was used to aid in writing code for the app. Gemini in Google Colab was utilized to help generate scripts for several figures based on Excel data. Grammarly was also used sparingly to correct sentence structure. No other artificial intelligence technologies were utilized in the process of conducting this study or manuscript writing.

### Human Testing

This study was conducted in accordance with the Institutional Review Board at UC Davis. The photos/videos of a participant was obtained voluntarily and taken by another participant. No other personal identifiers were kept or recorded from participants.

### Animal Usage

The cadaver head utilized for this study was provided by the UC Davis SVM’s Dentistry and Oral surgery Department and was sourced from euthanasia unrelated to this research.

### Software

*Project link:* https://github.com/ytipirneni/Vet_AR.git*. Operating system:* NebulaOS (Android 14), MacOS. *Programming language:* Unity (C#), Python. *Restrictions for non-academic use:* MIT License © 2025 Yash Tipirneni. The software may be used, modified, and distributed freely for academic and non-academic purposes.

### Data Availability Statement

Original datasets are available in a publicly accessible repository: https://doi.org/10.5061/dryad.n5tb2rc8t.

## Author Contributions

*Conceptualization:* YT, SG. *Funding acquisition:* YT, SG. *Resources:* YT, BA, SG. *Software:* YT. *Project administration:* YT, SG. *Supervision:* SG. *Methodology:* YT, SG. *Investigation:* YT. *Formal analysis:* YT, AB. *Visualization:* YT, CS. *Writing – original draft:* YT. *Writing – review & editing:* YT, AB, SG.

## References

1. Riggs J, Adams VJ, Hermer J v, Dobson JM, Murphy S, Ladlow JF. Outcomes following surgical excision or surgical excision combined with adjunctive, hypofractionated radiotherapy in dogs with oral squamous cell carcinoma or fibrosarcoma. Journal of the American Veterinary Medical Association. 2018;253(1):73–83.

2. Tuohy JL, Selmic LE, Worley DR, Ehrhart NP, Withrow SJ. Outcome following curative-intent surgery for oral melanoma in dogs: 70 cases (1998–2011). Journal of the American Veterinary Medical Association. 2014;245(11):1266–1273.

3. MacLellan RH, Rawlinson JE, Rao S, Worley DR. Intraoperative and postoperative complications of partial maxillectomy for the treatment of oral tumors in dogs. Journal of the American Veterinary Medical Association. 2018;252(12):1538–1547.

4. Sharma S, Boston SE, Skinner OT, et al. Survival time of juvenile dogs with oral squamous cell carcinoma treated with surgery alone: a Veterinary Society of Surgical Oncology retrospective study. Veterinary Surgery. 2021;50(4):740–747.

5. Milovancev M, Russell DS. Surgical margins in the veterinary cancer patient. Veterinary and comparative oncology. 2017;15(4):1136–1157.

6. Culp WTN, Ehrhart N, Withrow SJ, et al. Results of surgical excision and evaluation of factors associated with survival time in dogs with lingual neoplasia: 97 cases (1995–2008). Journal of the American Veterinary Medical Association. 2013;242(10):1392–1397.

7. Fransvea G, Moccia S, Bianchi F, et al. Intraoperative-technologies advancements in automated cancer detection: a narrative review. In: 2021 IEEE International Workshop on Metrology for Industry 4.0 & IoT (MetroInd4. 0&IoT). ieee; 2021:128–133.

8. Holt D, Okusanya O, Judy R, et al. Intraoperative near-infrared imaging can distinguish cancer from normal tissue but not inflammation. PloS one. 2014;9(7):e103342.

9. Alfonso Garcia A, Bec J, Weyers B, et al. Mesoscopic fluorescence lifetime imaging: Fundamental principles, clinical applications and future directions. Journal of biophotonics. 2021;14(6):e202000472.

10. Palmisciano P, Strong B, Strong E, Shahlaie K. Augmented Reality in Skull Base Oncology: A Pilot Study Focused on Tumor Resection and Skull Base Reconstruction. Journal of Neurological Surgery Part B: Skull Base. 2025;86(S 01):P359.

11. Sinha RY, Raje SR, Rao GA. Three-dimensional laparoscopy: principles and practice. Journal of minimal access surgery. 2017;13(3):165–169.

12. Blavier A, Nyssen AS. The effect of 2D and 3D visual modes on surgical task performance: role of expertise and adaptation processes. Cognition, technology & work. 2014;16(4):509–518.

13. Hatzipanayioti A, Bodenstedt S, von Bechtolsheim F, et al. Associations between binocular depth perception and performance gains in laparoscopic skill acquisition. Frontiers in Human Neuroscience. 2021;15:675700.

14. Beattie KL, Hill A, Horswill MS, Grove PM, Stevenson ARL. Laparoscopic skills training: the effects of viewing mode (2D vs. 3D) on skill acquisition and transfer. Surgical Endoscopy. 2021;35(8):4332–4344.

15. Vávra P, Roman J, Zonča P, et al. Recent Development of Augmented Reality in Surgery: A Review. Journal of Healthcare Engineering. 2017;2017. doi:10.1155/2017/4574172

16. Su Y xiong, Thieringer FM, Fernandes R, Parmar S. Virtual surgical planning and 3d printing in head and neck tumor resection and reconstruction. Frontiers in Oncology. 2022;12:960545.

17. Ahn D, Strong E, Marston A, et al. Clinical Application of Augmented Reality in Oral & Maxillofacial Surgery: A Multi-Center Case Series. International Journal of Oral and Maxillofacial Surgery. 2025;54:377.

18. Wu CC, Shih KT, Huang JW, Chen HH. A novel birdbath eyepiece for light field AR glasses. In: Optical Architectures for Displays and Sensing in Augmented, Virtual, and Mixed Reality (AR, VR, MR) IV. Vol 12449. SPIE; 2023:116–123.

19. Fida B, Cutolo F, di Franco G, Ferrari M, Ferrari V. Augmented reality in open surgery. Updates in surgery. 2018;70(3):389–400.

20. Aghapour M, Bockstahler B. State of the art and future prospects of virtual and augmented reality in veterinary medicine: a systematic review. Animals. 2022;12(24):3517.

21. XReal. Hand tracking: input and interactions. XReal Developer Docs. Published March 2025. Accessed 2025 Nov 15. Available from: https://docs.xreal.com/.

22. Unity Technologies. Unity user manual. Published 2025. Accessed 2025 Nov 15. Available from: https://docs.unity3d.com/.

23. Gechlik J, Sedrakyan H. Gauss’s Area Formula for Irregular Shapes. Ohio Journal of School Mathematics. 2024;97:12–29.

24. Fan YC, Zheng LJ, Liu YC. 3D environment measurement and reconstruction based on LiDAR. In: 2018 IEEE International Instrumentation and Measurement Technology Conference (I2MTC). IEEE; 2018:1–4.

25. Polycam Inc. *Polycam* [iOS application]. Published 2025. Accessed 2025 Nov 15. Available from: https://poly.cam.

26. İnal CB, Güngör MB, Nemli SK. Using a smartphone three dimensional scanning application (Polycam) to three dimensionally print an ear cast: A technique. The Journal of prosthetic dentistry. Published online 2023.

27. Wagner A, Undt G, Watzinger F, et al. Principles of computer-assisted arthroscopy of the temporomandibular joint with optoelectronic tracking technology. *Oral Surgery, Oral Medicine, Oral Pathology*, Oral Radiology, and Endodontology. 2001;92(1):30–37.

28. Wilkat M, Kübler N, Rana M. Advances in the resection and reconstruction of midfacial tumors through computer assisted surgery. Frontiers in Oncology. 2021;11:719528.

29. Strong EB, Patel A, Marston AP, et al. Augmented Reality Navigation in Craniomaxillofacial/Head and Neck Surgery. OTO open. 2025;9(2):e70108.

30. Devaraji M. The role of neurogenomics in personalized neurosurgical interventions: a new frontier in precision medicine. Neurosurgical Review. 2024;47(1):488.

31. Wanzel KR, Hamstra SJ, Caminiti MF, Anastakis DJ, Grober ED, Reznick RK. Visual-spatial ability correlates with efficiency of hand motion and successful surgical performance. Surgery. 2003;134(5):750–757.

32. Koulieris GA, Akşit K, Stengel M, Mantiuk RK, Mania K, Richardt C. Near eye display and tracking technologies for virtual and augmented reality. In: Computer Graphics Forum. Vol 38. Wiley Online Library; 2019:493–519.

