## Supplementary material for "Assessment of Augmented Reality Glasses for Spatial Tracking and Intraoperative Annotation in Veterinary Surgery": Figure

**Figures**

1. Three panels side by side. The left panel shows a hand without any annotations. The middle panel shows the same hand and a vector-line overlay with white lines and red dots. The right panel shows the vector-line overlay on a black background.
2. Two panels side by side. The left panel shows a photo of a pitbull head with a clay tumor mold on the top teeth, a mouth piece between the left canines, and metal lip retractors pulling back the cheeks. The background is a photo of a blood-stained paper towel in black-and-white. There is a marker that says 0.04 m next to the mouthpiece. The left panel shows a 3D reconstruction of the same head on a solid-colored background. There is a marker that says 0.04 m on the mouthpiece.
3. Three side by side panels showing the three AR testing steps. The left panel is titled 1 - practice, and a finger is touching a silhouette of a dog head while drawing points with a 2 minute time in the background. The middle panel is titled 2 - coordinates. The top half titled transfer distance has a computer screen with a dot on the dog head and a 2 second timer, and a finger is placed next to the same dog head without that dot. The bottom half is titled direct distance, and has a finger placed next to the same dog head with the same point already visible. The right panel is titled 3 - ROI. It is set up almost exactly as the middle panel, except the monitor has a 5 second timer, the labels are area instead of distance, and the dots are replaced by an oblong region.
4. Three panels side by side. The left panel shows a side view of the dog head with the locations of 5 points. The middle panel shows the average distance error in millimeters for the 5 transfer points on the left side and direct points on the right side for all participants. Each group has a diamond-shaped average bar with standard deviation lines, and grey lines connect transfer and direct data for each participant. The right panel is set up almost similarly, except the units on the y-axis are time in seconds.
5. Three panels side by side. The left panel shows a side view of the dog head with the locations of 3 regions. The middle panel is a graph comparing non-overlap percent and region coverage percent on the x and y-axis, respectively, for each participant. A legend shows that circles correspond to transfer data while the squares represent direct data. The right panel shows the average time taken in seconds by each participant for plotting the regions. A grey line connects the transfer data on the left side to the direct data on the right side for each participant.
6. Four panels are grouped in a 2 by 2 format. All panels have distance error on the y-axis. The top left panel presents the distance error data sorted from least to greatest and color coded based on specialty. The top right panel has the years of experience on the x-axis, and a trendline with a negative slope is shown. The bottom left panel compares Rx data with Rx data side by side with corresponding average and standard deviation bars. The bottom right panel has the arm-length in centimeters on the x-axis, and a trendline with a slope close to zero is shown.
7. The survey results for 10 different questions are displayed as horizontal stacked bar graphs. The results are color coded and displayed as percentages of the categories: strongly disagree, disagree, neutral, agree, and strongly disagree.
8. Two side by side illustrations of a surgeon holding a scalpel. The left panel titled camera and monitor setup has the surgeon looking up at a television screen that shows a classifier overlay on the surgical field of the canine oral tumor. The right panel titled AR headset has the surgeon wearing a headset looking directly at the patient. A callout box shows the view from the headset where they can see the same classifier overlay projected directly on the surgical field.

**Supplemental material**

1. A participant who is sitting down and wearing AR glasses interacts with a dog head hologram by drawing virtual annotations on its upper lip. A vector-line overlay is also visible on their right hand. A cable extends down from the glasses and connects to a phone on the table.
2. Scatter plots showing distance errors in millimeters for each of the five coordinates, with separate categories for transfer and direct conditions.
3. Scatter plots showing task completion times in seconds for each coordinate, for the transfer and direct groups.
4. Scatter plots showing the relationship between ROI coverage percentage and non-overlap percentage for each ROI, with separate points for transfer and direct conditions.
5. Scatter plots showing task completion times in seconds for each of the 3 ROIs, for the transfer and direct groups.
6. Two plots of years of experience versus normalized distance error in millimeters. The scatter plot’s line of best fit has a negative slope. The box-and-whisker compared residents and specialists.
