## Supplementary figures and images for "Assessment of Augmented Reality Glasses for Spatial Tracking and Intraoperative Annotation in Veterinary Surgery"

### Dog_Scan.jpg

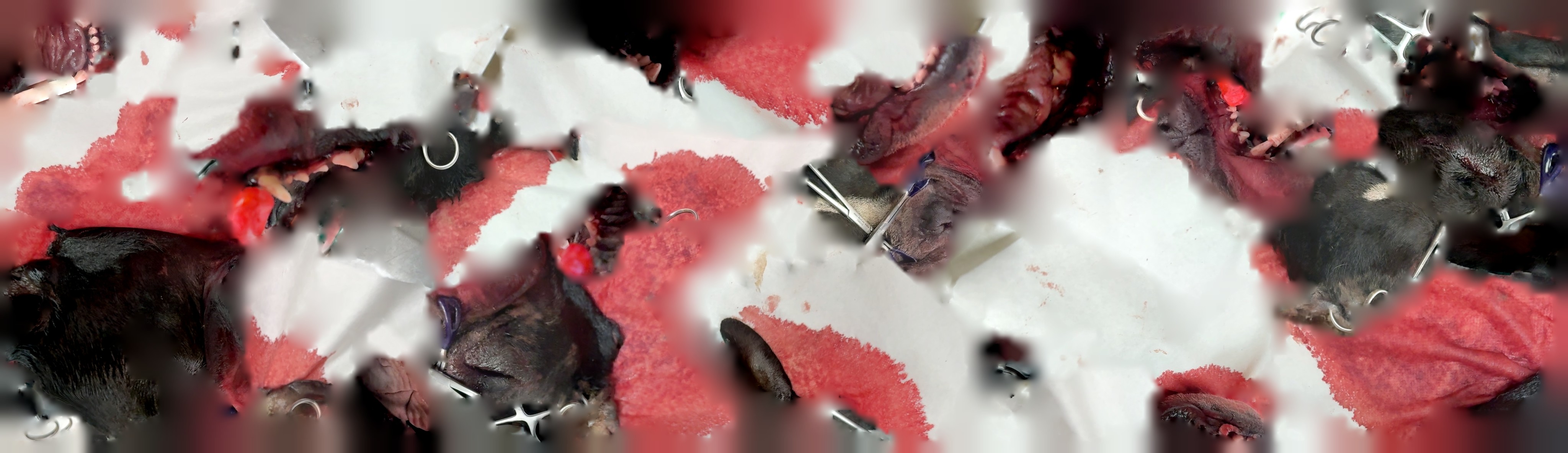
