## Supplementary Material for "Assessment of Augmented Reality Glasses for Spatial Tracking and Intraoperative Annotation in Veterinary Surgery"

*Supplementary Materials*

### Supplementary Material

#### Supplementary Code:

The full source code for the AR application is available at: <https://github.com/ytipirneni/Vet_AR.git> (Accessed November 27, 2025).

Operating system: NebulaOS (Android 14), macOS.

Programming language: Unity (C#), Python.

License: MIT License © 2025 Yash Tipirneni. The software is open-source and may be used, modified, and distributed freely for academic and non-academic purposes.

#### Supplementary 3-D File:

See *Dog_Scan.zip* for the 3-D model (.OBJ) of the dog head.

#### Supplementary Data:

The original dataset is available in a publicly accessible repository: <https://doi.org/10.5061/dryad.n5tb2rc8t>.

### Supplementary Videos

*Supplementary Video S1*: Demonstration of the practice step, which had a 2-minute limit.

*Supplementary Video S2*: Demonstration of the coordinate step for both transfer and direct tasks.

*Supplementary Video S1:* Demonstration of the outline steps for both transfer and direct tasks.

### Supplementary Figures

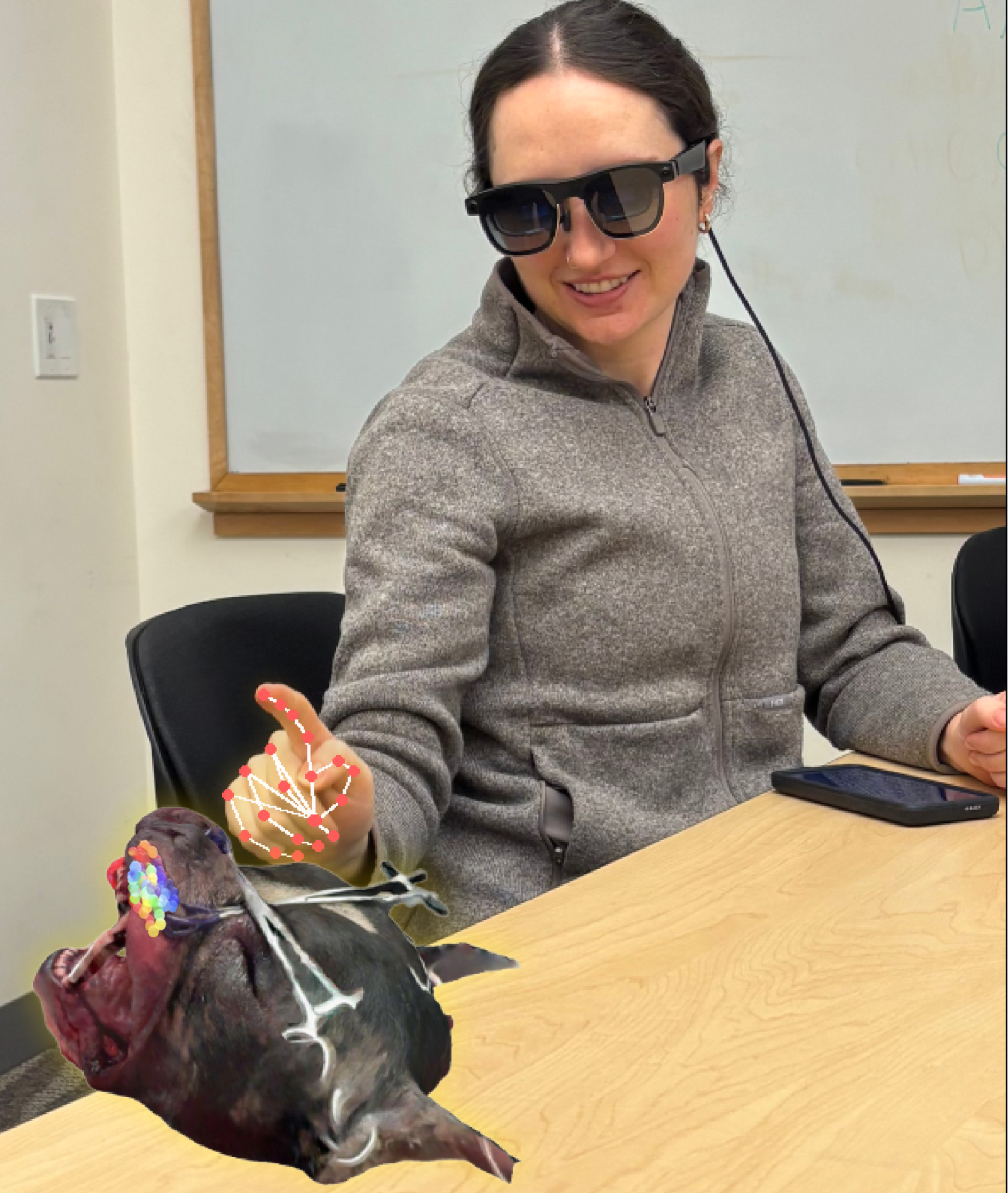

**Supplementary Figure S1.** Simulation of what a study participant sees when they are testing the program. The participant is practicing drawing on the virtual dog head with their pointer finger. The augmented reality (AR) glasses are connected to a controller with a USB-C cable.

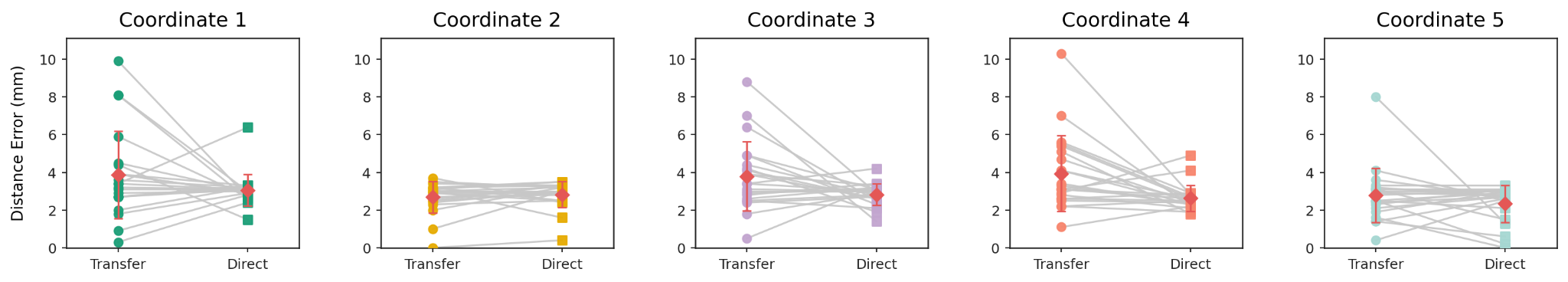
**Supplementary Figure S2.** Comparison of the distance error for transfer vs direct tasks for each of the 5 coordinates. Each plot follows a similar trend to Figure 3B and is color-coded the same as the targets in Figure 3A.

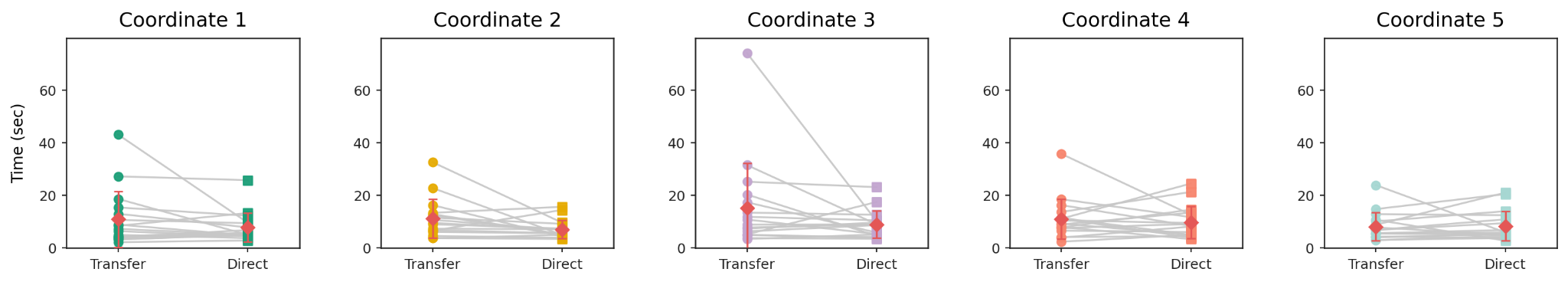
**Supplementary Figure S3.** Comparison of the time taken for transfer vs direct groups for each of the 5 coordinates. Each plot follows a similar trend to Figure 3C and is color-coded the same as the targets in Figure 3A.

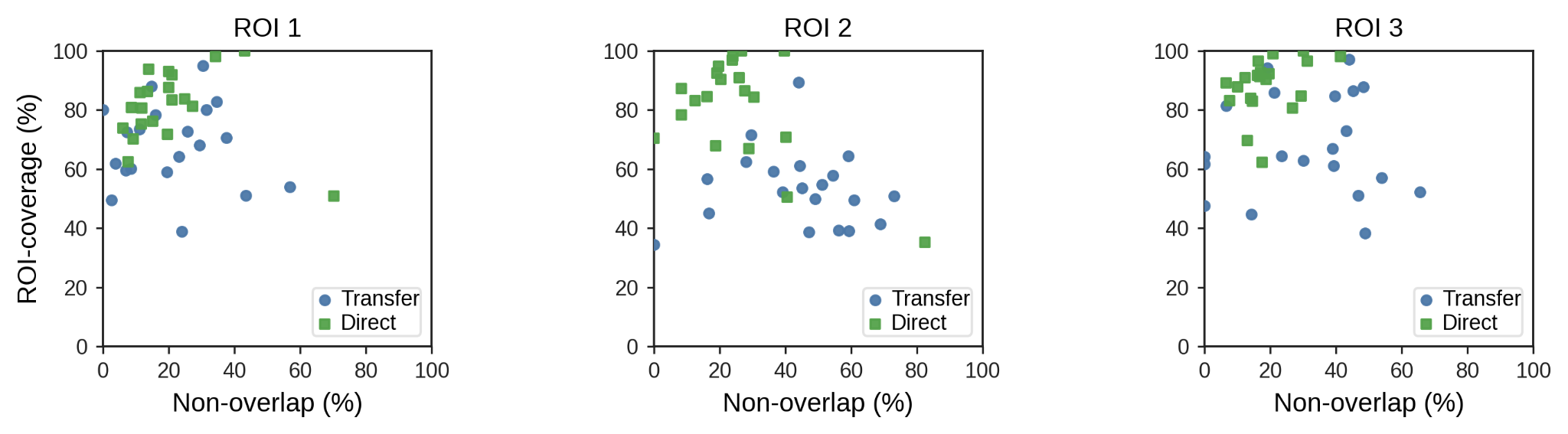
**Supplementary Figure S4.** Comparison of non-overlap percentage with region of interest (ROI) coverage for the transfer and direct groups. Each plot corresponds to a different target ROI from Figure 3D, and the data follows a similar distribution to Figure 3E.

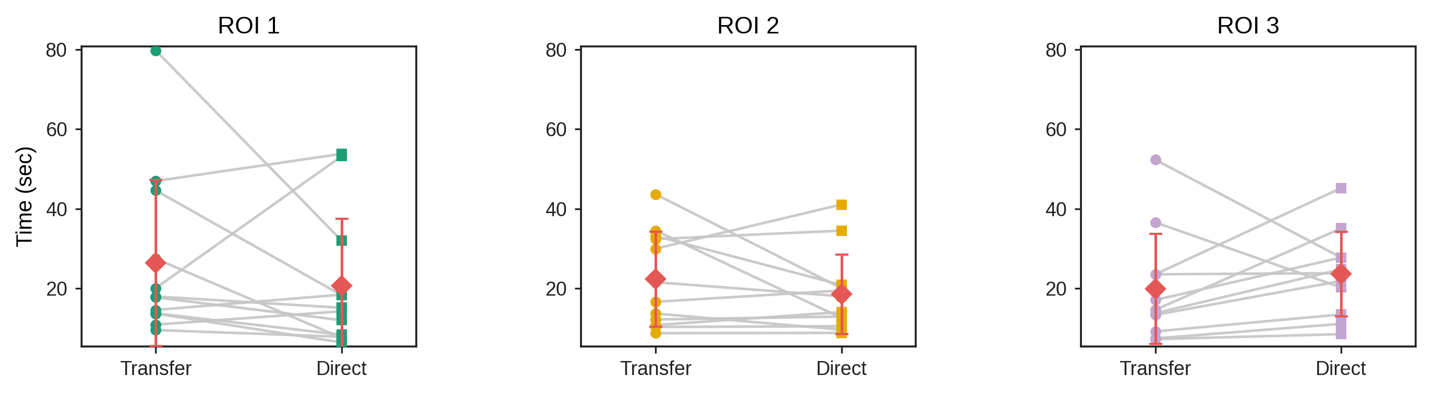
**Supplementary Figure S5.** Comparison of the time taken for the transfer and direct groups with each of the regions of interest (ROI). Each plot is color-coded the same as the ROIs in Figure 3D, and the data follows a similar distribution to Figure 3F.

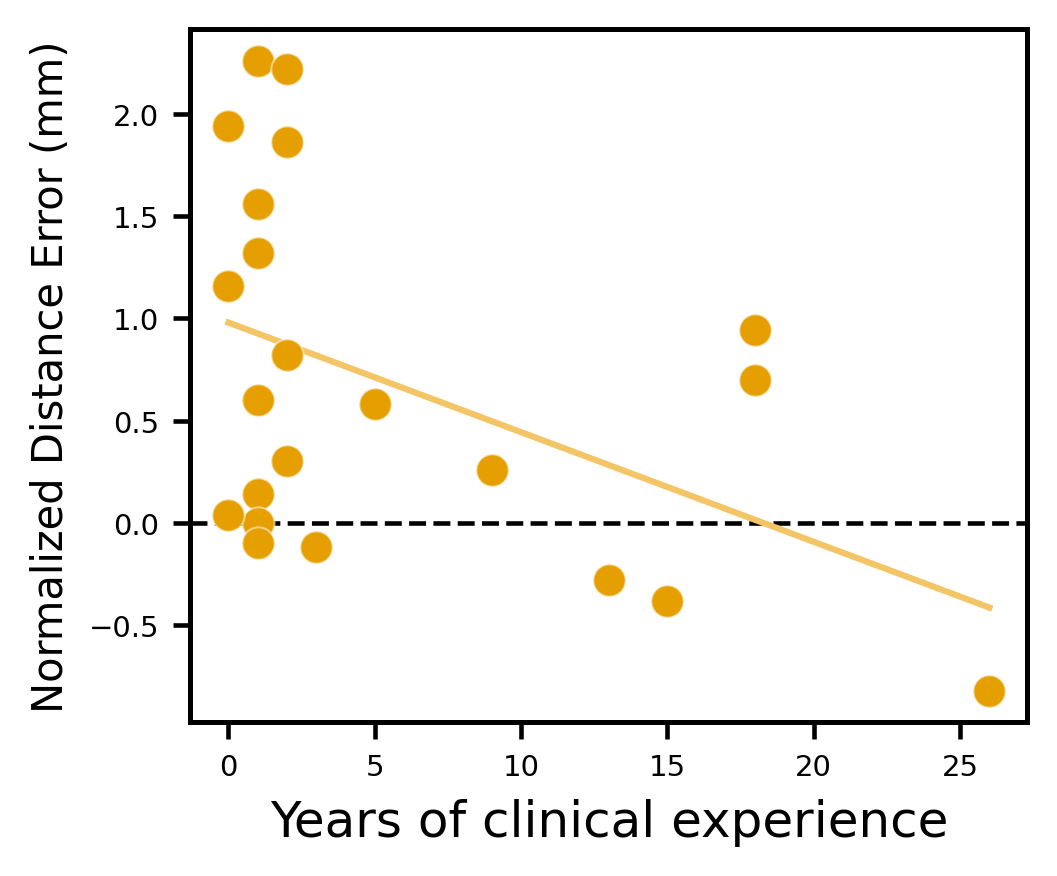

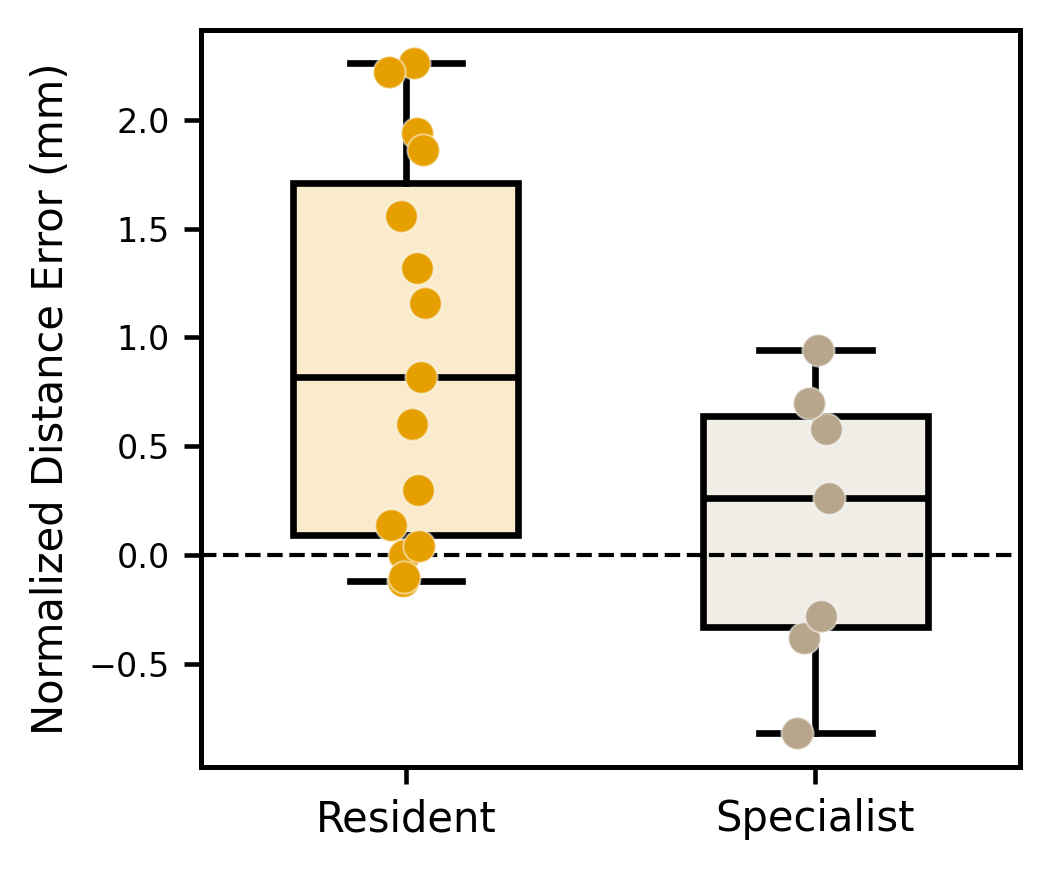

**Supplementary Figure S6.** Comparison of years of clinical experience with the normalized distance error in millimeters similar to Figure 4B. The left plot is a scatter plot with a linear regression of *-0.0536x + 0.979 (R^2^ = 0.207).* The right plot is a box-and-whisker plot of residents (0-3 years) and specialists (4+ years). For the box-and-whisker plot, the solid line within the box represents the median, the lower and upper limits of the box represent the first and third quartile, respectively, and the whiskers delimit the range.
